# Development of an objective index, neural activity score (NAS), reveals neural network ontogeny and treatment effects on microelectrode arrays

**DOI:** 10.1101/2020.07.21.213629

**Authors:** Austin P. Passaro, Onur Aydin, M. Taher A. Saif, Steven L. Stice

## Abstract

Microelectrode arrays (MEAs) are valuable tools for electrophysiological analysis at a cellular population level, providing assessment of neural network health and development. Analysis can be complex, however, requiring intensive processing of large high-dimensional data sets consisting of many activity parameters. As a result, valuable information is lost, as studies subjectively report relatively few metrics in the interest of simplicity and clarity.

From a screening perspective, many groups report simple overall activity; we are more interested in culture health and changes in network connectivity that may not be evident from basic activity parameters. For example, general changes in overall firing rate – the most commonly reported parameter – provide no information on network development or burst character, which could change independently. Our goal was to develop a fast objective process to capture most, if not all, the valuable information gained when using MEAs in neural development and toxicity studies.

We implemented principal component analysis (PCA) to reduce the high dimensionality of MEA data. Upon analysis, we found that the first principal component was strongly correlated to time, representing neural culture development; therefore, factor loadings were used to create a single index score – named neural activity score (NAS) – reflective of neural maturation. To validate this score, we applied it to studies analyzing various treatments. In all cases, NAS accurately recapitulated expected results, suggesting this method is viable. This approach may be improved with larger training data sets and can be shared with other researchers using MEAs to analyze complicated treatment effects and multicellular interactions.

**Author Summary:** Analyzing neural activity has important applications such as basic neuroscience research, understanding neurological diseases, drug development, and toxicity screening. Technology for recording neural activity continues to develop, producing large data sets that provide complex information about neuronal function. One specific technology, microelectrode arrays (MEAs), has recently given researchers the ability to record developing neural networks with potential to provide valuable insight into developmental processes and pathological conditions. However, the complex data generated by these systems can be challenging to analyze objectively and quantitatively, hindering the potential of MEAs, especially for high-throughput approaches, such as drug development and toxicity screening, which require quick, simple, and accurate quantification. Therefore, we have developed an index for simple quantification and evaluation of neural network maturation and the effects of perturbation. We present validation of our approach using several treatments and culture conditions, as well as a meta-analysis of toxicological screening data to compare our approach to current methods. In addition to providing a simple quantification method for neural network activity in various conditions, our method provides potential for improved results interpretation in toxicity screening and drug development.

## Introduction

Micro- (or multi-)electrode arrays (MEAs) are valuable tools for network-level electrophysiological analysis of neuronal populations [1–4]. While sacrificing single cell resolution compared to traditional patch clamp electrophysiology, MEAs allow for recordings of entire neural networks both *in vitro* and *in vivo* and can be used to study dynamic network properties and development, either spontaneously or in response to stimulation or treatment. During recording, action potentials, or spikes, are detected *via* recording the corresponding voltage changes in the extracellular environment. Analysis of spike patterns provides network characteristics such as firing rate and network synchrony (see S1 Table) for list of all measured parameters), which are useful when determining neuronal network function and/or response to perturbation *(i.e.* stimulation or pharmacological treatment) [5,6].

Given these advantages, along with the advent of multi-well MEA plates that allow for higher-throughput screening and more complex experimental design, MEAs have seen widespread adoption from characterizing neural maturation to toxicity screening and drug development. Interestingly, despite the adoption of MEAs for these screening approaches, analysis has typically been limited to mean firing rate and other metrics of overall activity [7,8]. This limited analysis severely underutilizes MEA capabilities and may result in “false-negative” screening results, as only conditions or compounds that increase or decrease overall neural activity will be registered as hits with no regard to other aspects of neural network functionality or ontogeny. Current MEA analysis methods require the use of raster plots to visualize network development or individual parameter analysis, which are qualitative and difficult to interpret, respectively. While a general pattern of network development from sporadic spikes to sporadic bursts to coordinated synchronous network bursts has been well-described in previous studies [2,3,9,10], there is currently a lack of sufficient methods to quantify this observed ontogeny.

Here, we developed a method implementing dimensionality reduction techniques, specifically principal component analysis (PCA), to create a singular index score – named neural activity score (NAS) – reflective of neural network ontogeny. NAS serves as an easily interpretable measurement to evaluate spontaneous network development in simple and complex cultures *(i.e.* neuron-glia co-cultures) or effects of various treatments (*i.e.* soluble factors or stimulation). We present validation of this method in several experiments, including a culture media comparison, various conditioned media treatments, and a microglia-neuron co-culture system, demonstrating the ability to measure both positive and negative effects on neural network activity and further interrogate toxicological screening, evaluating sensitivity on potential toxic compounds.

## Results

### Neural network ontogeny revealed by microelectrode array

Mouse embryonic stem cells were cultured and differentiated, resulting in cultures containing a mixture of motor neurons, excitatory and inhibitory neurons, and glial cells [11]. These neural cultures were allowed to mature on 48-well MEA plates over a 19-day period, typical for neuron maturation and network formation for these cells [12]. Activity was detected at approximately 5 days post-seeding (days *in vitro;* DIV) and increased gradually until reaching a plateau over the last several days of recording (16-19 DIV). Qualitatively, raster plots generated at various time points throughout the recording period demonstrate an expected pattern of network development: sparse and sporadic spikes appearing first, followed by sporadic bursts, followed eventually by synchronous network bursts (Fig 1A-F). While this emergent development is evident from the raster plots, it is difficult to quantify. Quantification of several spike, burst, and network/synchrony metrics reveals general increases over time in these categories (Fig 1G-N), but current MEA analysis methods do not allow for simple quantification of network ontogeny incorporating these and other activity metrics.

**Fig 1.**
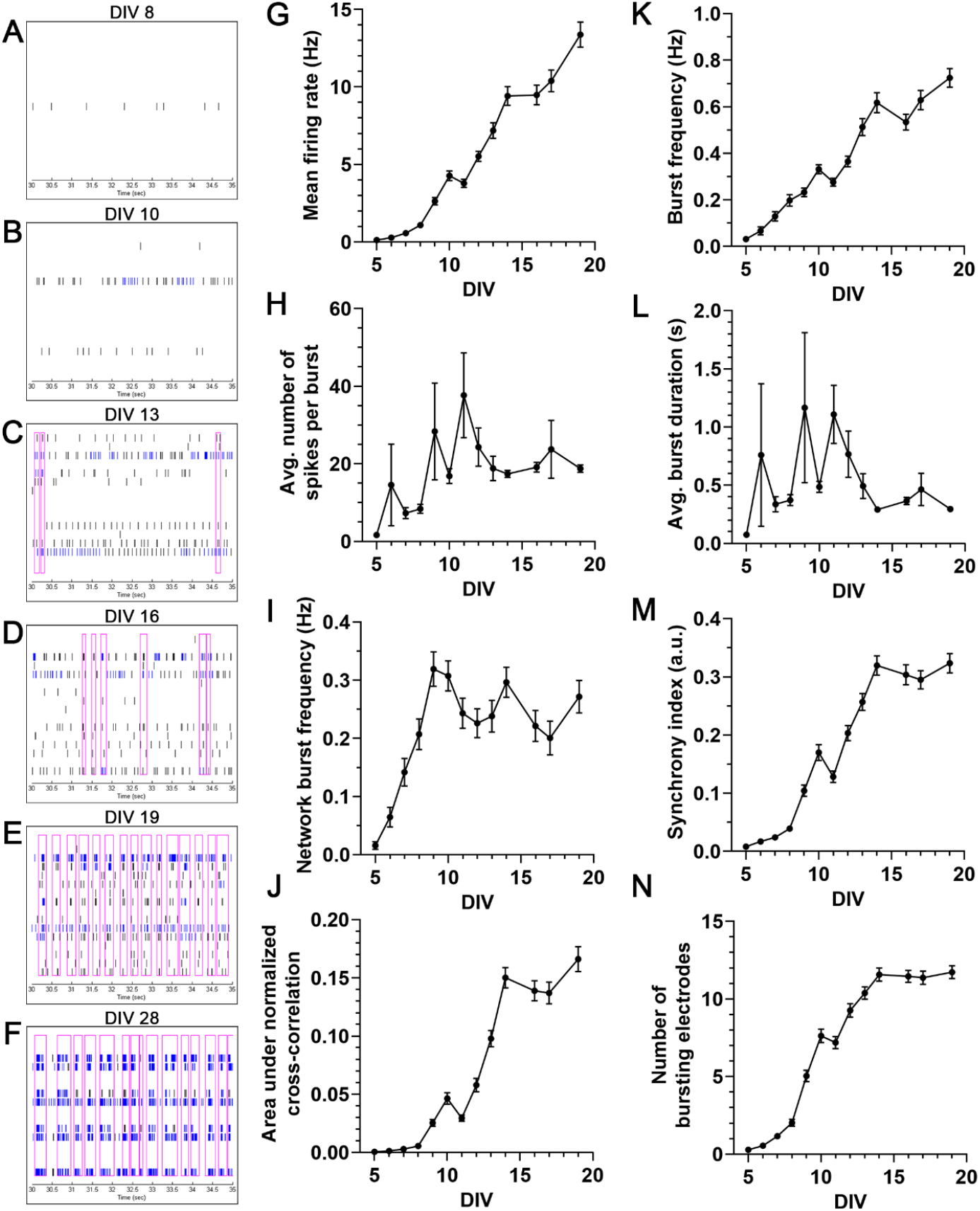
Neural network ontogeny revealed by microelectrode array. (A-F) Representative raster plots of one well over time, demonstrating qualitative network development. Each plot is 5 seconds for sufficient spike and burst resolution, and horizontal rows correspond to one channel/electrode, each. Note the changes over time: few spikes on few channels (DIV 8) to more spikes on more channels (DIV 10) to sporadic bursts (DIV 13, 16) to rhythmic network bursts (DIV 19) to stronger, rhythmic network bursts (DIV 28). (G-N) Line graphs of 8 example individual MEA parameters covering major categories (activity, bursting, network bursting, synchrony).

### Principal component analysis of MEA parameters reveals temporal correlation, allowing for neural activity score derivation

Given the complexity and multivariate nature of the data, PCA was performed to reduce dimensionality and allow for easier visualization. After standard score normalization, all of the aforementioned parameters at all time points were included as data points for PCA. Examining the two-dimensional projection of the first two principal components revealed a distinct pattern in the data (Fig 2A). Adding a dimension of time (*via* colormap), this pattern was revealed to be a temporal separation of the data points, especially along the first principal component. Statistically, linear regression analysis supported this temporal component, as principal component 1 (PC1) is strongly correlated to time (Fig 2B; R^2^=0.5441, p<0.0001), indicating recapitulation of network ontogeny and maturation. After confirmation of this relationship, factor loading values for PC1 were examined to determine which factors (MEA parameters) contributed most strongly to this component. While substantial contributions were observed for many parameters, the strongest metrics were burst percentage, network burst percentage, number of spikes per burst, number of bursting electrodes, number of spikes per network burst, and synchrony index (Table 1). Notably, mean firing rate, the most common parameter analyzed in MEA studies, was the 11^th^-strongest contributor. Finally, these factor loading values were used to develop an individual index score – NAS (Equation 1; see Methods). As NAS represents all aspects of neural network activity, it allows for assessment of neuronal network ontogeny and evaluation of the effects of various perturbations, such as stimulation, pharmacological treatment, or alternative culture conditions, which can typically be difficult to analyze if various parameters do not exhibit unidirectional changes. Additionally, NAS reduces the high variation often observed in individual MEA parameters, as evidenced by lower coefficient of variation for 24/25 (96%) measured parameters (S1 Fig).

**Fig 2.**
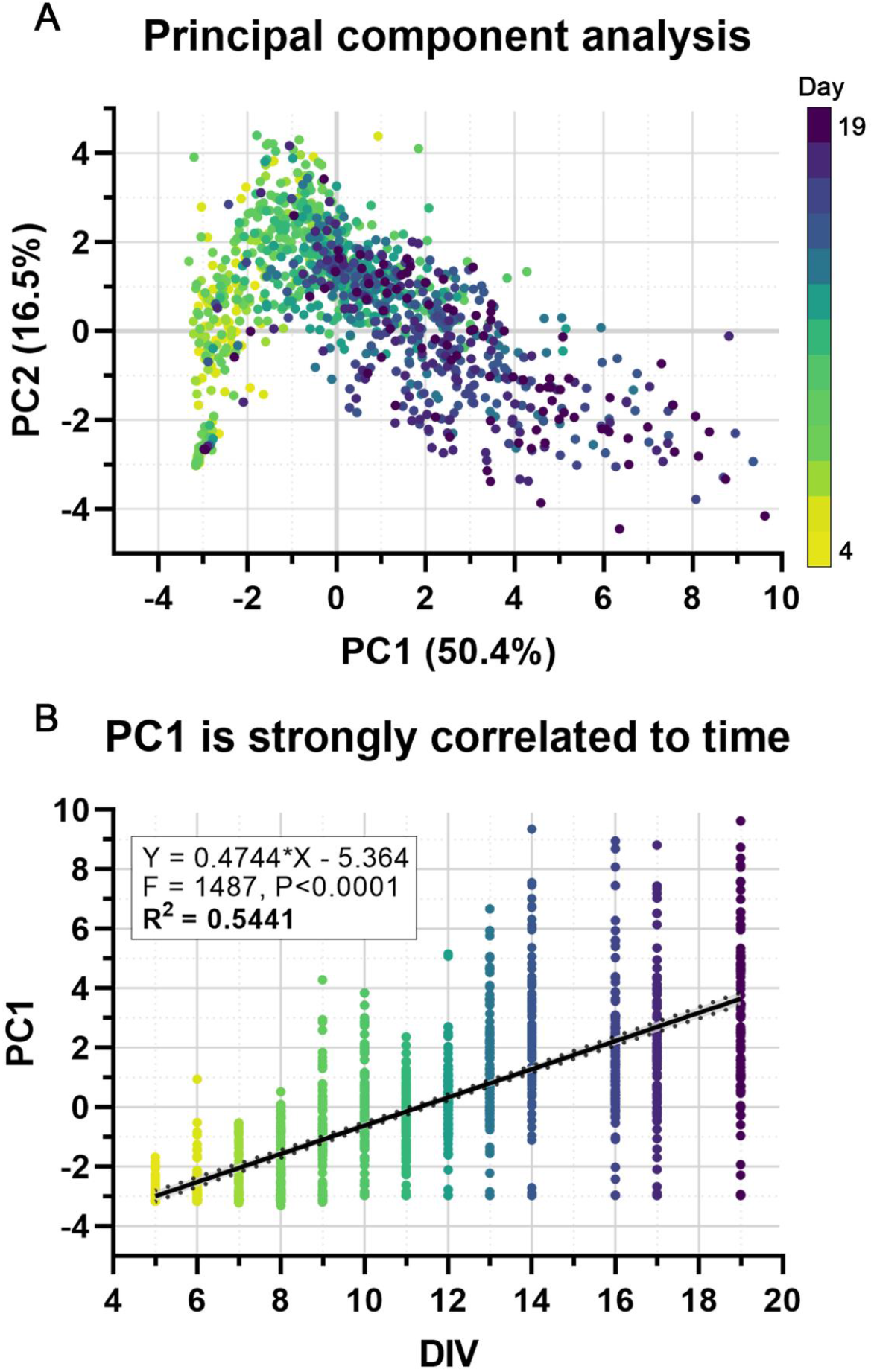
Principal component analysis of MEA parameters reveals temporal correlation. (A) The first two principal components (accounting for 66.9% of total variation), colored by time (yellow > green > blue > purple), showing a distinct pattern of separation/progression. (B) Principal component 1 (PC1) is positively correlated with time. Linear regression analysis confirms this strong correlation (R^2^ = 0.5541, F = 1487, p<0.0001).

**Table 1.** Factor loading values for principal components 1-5 for all MEA parameters analyzed.

| *Parameter* | *PC1* | *PC2* | *PC3* | *PC4* | *PC5* |
| --- | --- | --- | --- | --- | --- |
| Burst Percentage - Avg | 0.938953 | -0.21655 | -0.09068 | 0.095365 | -0.03905 |
| Network Burst Percentage | 0.930335 | -0.16411 | -0.03479 | -0.00912 | 0.030603 |
| Number of Spikes per Burst - Avg | 0.927633 | -0.07813 | 0.131878 | 0.180872 | -0.20604 |
| Number of Bursting Electrodes | 0.924737 | 0.11382 | -0.13932 | -0.05683 | -0.03851 |
| Number of Spikes per Network Burst per Channel - Avg | 0.916818 | -0.09696 | 0.059892 | -0.0695 | -0.30286 |
| Synchrony Index | 0.91299 | -0.25337 | -0.09404 | 0.126219 | -0.03106 |
| Number of Spikes per Network Burst - Avg | 0.90668 | -0.14074 | 0.063698 | -0.03136 | -0.3157 |
| ISI Coefficient of Variation - Avg | 0.875459 | -0.13787 | -0.14411 | 0.203831 | -0.09526 |
| Area Under Normalized Cross-Correlation | 0.866748 | -0.32439 | -0.0838 | 0.14749 | -0.01187 |
| Number of Elecs Participating in Burst - Avg | 0.866593 | 0.127533 | -0.16016 | -0.10207 | -0.04436 |
| Mean Firing Rate (Hz) | 0.787233 | -0.09261 | 0.483835 | -0.01736 | 0.111684 |
| Network IBI Coefficient of Variation | 0.739427 | 0.22799 | -0.36458 | -0.23674 | 0.12618 |
| Burst Duration - Avg (s) | 0.730618 | 0.521412 | 0.032794 | 0.197191 | -0.12086 |
| Normalized Duration IQR - Avg | 0.707402 | 0.327012 | -0.30471 | 0.026697 | 0.337296 |
| IBI Coefficient of Variation - Avg | 0.64845 | 0.459111 | -0.30775 | 0.114136 | 0.20907 |
| Network Normalized Duration IQR | 0.613893 | 0.064961 | -0.38534 | -0.17934 | 0.144007 |
| Area Under Cross-Correlation | 0.606648 | -0.40216 | 0.517657 | 0.237669 | -0.14355 |
| Burst Frequency - Avg (Hz) | 0.592944 | -0.04631 | 0.437397 | 0.034638 | 0.423479 |
| Network Burst Frequency (Hz) | 0.506543 | -0.0198 | 0.364146 | -0.01999 | 0.590348 |
| Network Burst Duration - Avg (sec) | 0.490791 | 0.412156 | 0.078862 | -0.54881 | -0.07344 |
| Width at Half Height of Cross-Correlation | 0.233083 | 0.626106 | 0.385101 | -0.42764 | -0.13819 |
| Width at Half Height of Normalized Cross-Correlation | 0.175127 | 0.727927 | 0.205821 | -0.28793 | -0.18294 |
| Mean ISI within Burst – Avg | 0.029257 | 0.891807 | 0.112554 | 0.360375 | 0.016162 |
| Median ISI within Burst - Avg | -0.05205 | 0.881542 | 0.1378 | 0.359244 | 0.017994 |
| Inter-Burst Interval - Avg (s) | -0.27591 | 0.571143 | -0.11858 | 0.370944 | -0.15048 |

Parameters sorted by descending PC1 loading value.

### Enhancement of neural network ontogeny is easily quantified via NAS

Regression analysis served as initial support that NAS accurately measures neural network ontogeny, but we also sought to experimentally validate NAS in several conditions to further confirm this recapitulation. For initial validation, several experiments were performed to analyze enhanced neural network ontogeny and activity in response to different conditions known to enhance neural activity – namely, optimized culture media [13] and muscle-conditioned media treatment [14]. To examine the effects of optimized culture media, mixed neural cultures (HBG3-derived) were grown on MEAs in two different media conditions: DMEM/F12 & Neurobasal-based medium (DMNB) or BrainPhys^TM^-based medium (BP). While DMNB has traditionally been widely used to culture HBG3-derived and other neural cell lines, BP was developed for electrophysiological applications due to a more physiologically relevant formulation, resulting in increased electrophysiological function of various cell lines [13]. However, BP has not been evaluated on HBG3-derived neural cultures. In both DMNB and BP groups, the neurons began showing activity at approximately day 5, increasing over 3 weeks, as expected; however, the cells cultured in BP exhibited enhanced activity and network development, as indicated by the significantly higher NAS (Fig 3A; p<0.0001, two-way repeated measures ANOVA).

**Fig 3.**
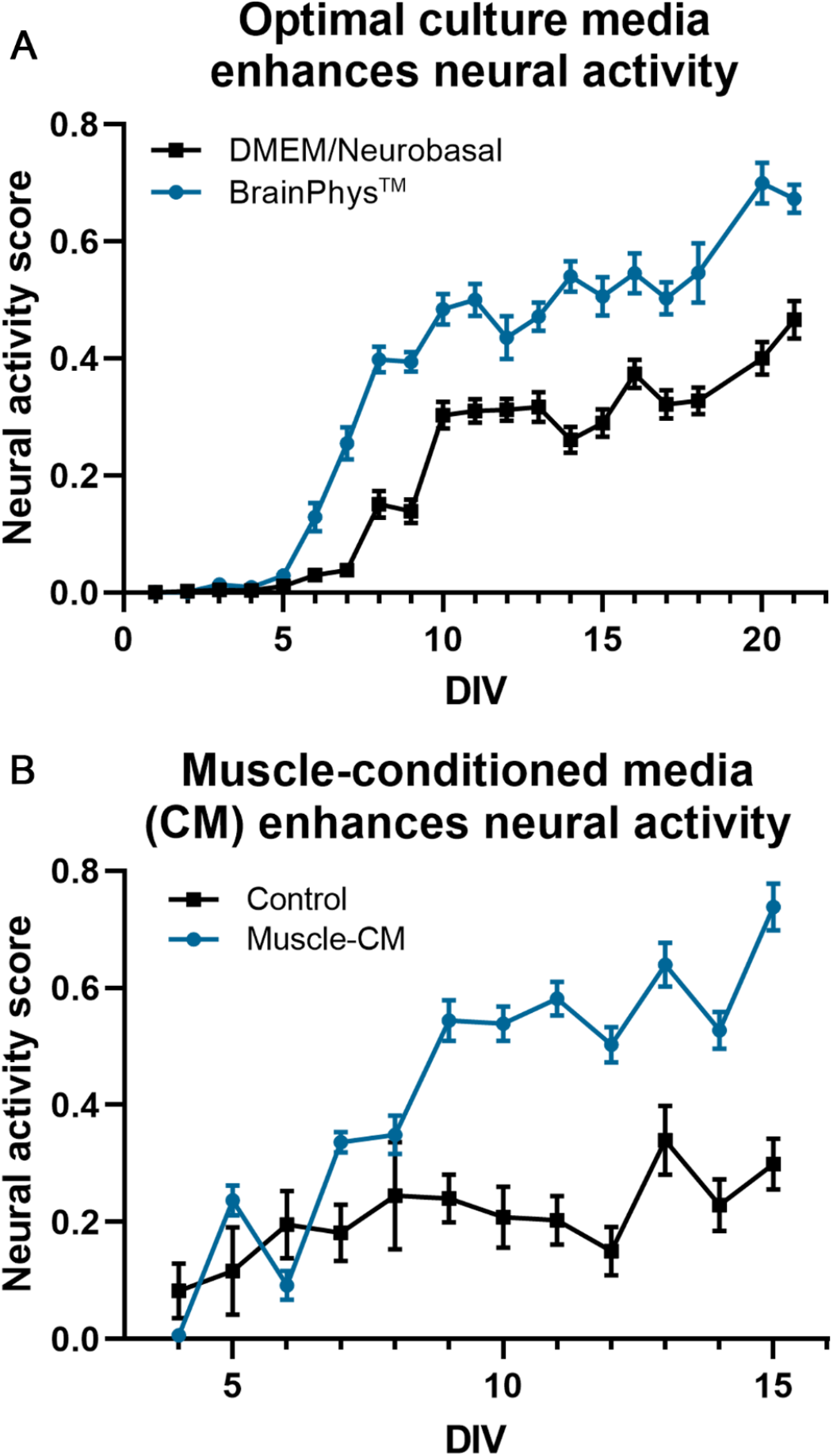
Enhancement of neural network ontogeny is easily quantified using neural activity score. (A) BrainPhys^TM^-based culture media results in clear enhancement of neural activity compared to traditional DMEM/Neurobasal-based media, and this enhancement is quantifiable via NAS (p<0.0001, two-way repeated measures ANOVA, n=24/group). (B) Muscle-conditioned media treatment results in similar enhancement of neural activity (p<0.0001, two-way repeated measures ANOVA, n=12/group).

To examine the effects of conditioned media on network ontogeny, mixed neural cells were treated with muscle cell (C2C12)-conditioned media (CM), which has previously been shown to significantly accelerate network activity and development [14]. Likewise, NAS analysis showed similar results and provided simple quantification (Fig 3B; p<0.0001, two-way repeated measures ANOVA) of this accelerated network development.

### Disruption of neural network ontogeny is easily quantified via NAS

In addition to measuring neural network activity enhancement, we also sought to validate NAS on more complex culture conditions and for quantifying disruption of network activity. Microglia, the resident immune cells of the central nervous system (CNS), are being increasingly implicated in neurodegenerative diseases and have been shown to be neurotoxic in many conditions [15–18]; therefore, we decided to explore co-culturing microglia with mixed neural cultures on MEAs. After allowing neurons to become active over 10 days, BV2 cells, an immortalized mouse microglia cell line, were added to the cultures at 8 different cell densities. We observed rapid disruption in network function in a clear cell density-dependent manner, with higher numbers of microglia relative to the neuronal population resulting in accelerated network disruption, as indicated by a decrease in NAS (Fig 4A; p<0.0001, twoway repeated measures ANOVA, Tukey’s post-hoc test).

**Fig 4.**
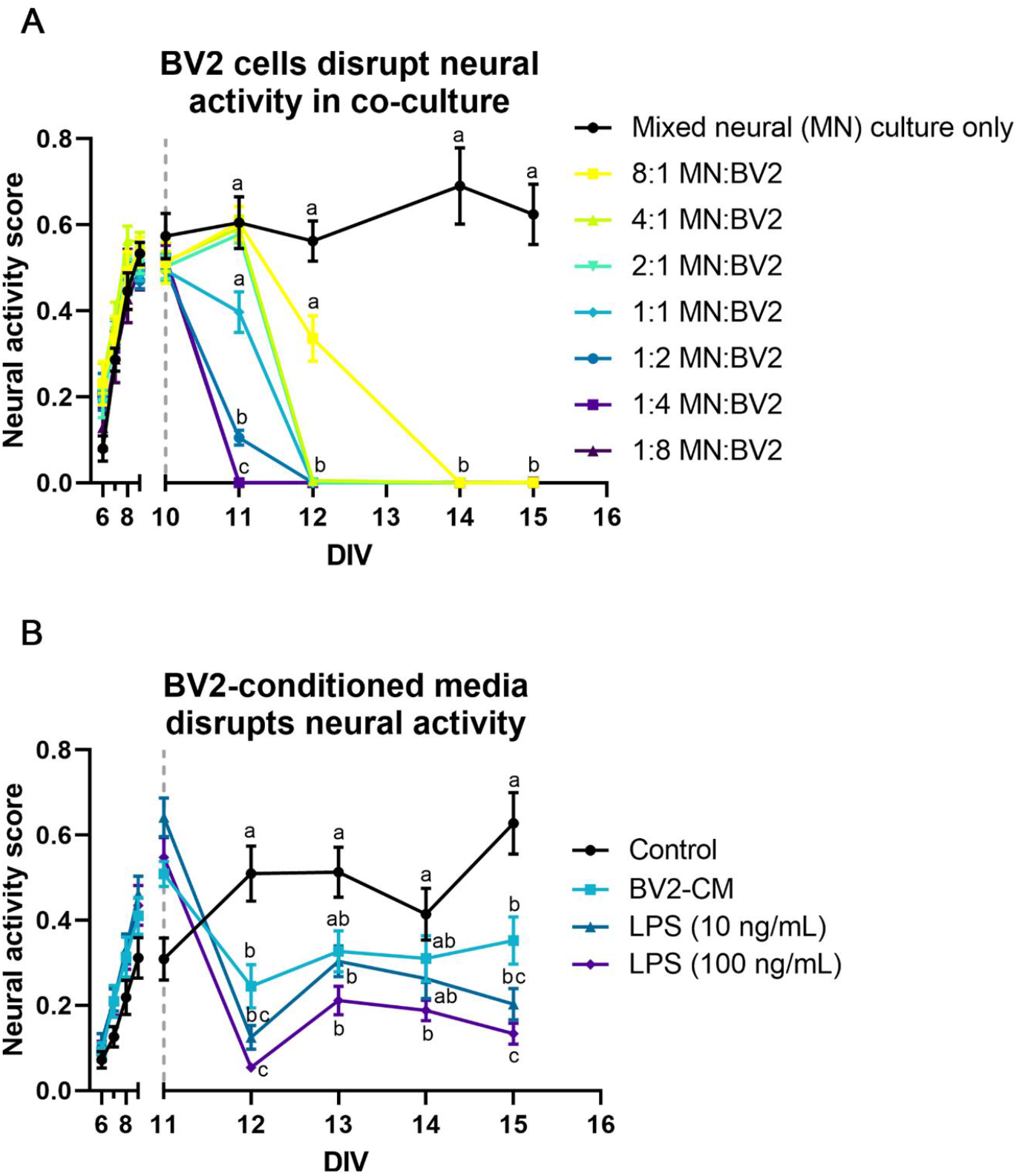
Disruption of neural network ontogeny is easily quantified using neural activity score. (A) Co-culturing mixed neural cultures and microglia (BV2 cells) results in a microglia concentrationdependent disruption of neural activity (p<0.0001, two-way repeated measures ANOVA, Tukey’s post-hoc test, n=6/group). (B) Similarly, BV2-conditioned media treatment resulted in a similar decrease (p<0.0193, two-way mixed ANOVA, Tukey’s post-hoc test). Additionally, 24-hour LPS treatment of BV2s prior to conditioned media collection exacerbated this disruption in a concentration-dependent manner (10 ng/mL, p=0.0003; 100 ng/mL, p<0.0001) (n=14/group except media control group, for which n=12). Grey dashed lines indicate time of BV2 or CM addition. Reported statistics are Tukey’s post-hoc comparisons 24 hours post-addition. Connecting letters on graphs indicate comparisons for other time points.

To examine whether this disruption is contact-dependent or the result of secreted factors, neural cultures were treated with BV2-conditioned media at 10 days (similarly to the co-culture experiment described above). Similar to the co-culture condition, BV2-conditioned media treatment also disrupted network function (Fig 4B), suggesting a role for microglia-secreted factors in neural network disruption. To examine whether this effect was exacerbated by microglial activation, BV2 cells were stimulated with two concentrations of the pro-inflammatory endotoxin lipopolysaccharide (LPS; 10 ng/mL and 100 ng/mL) for 24 hours prior to conditioned media collection. LPS-stimulated BV2-conditioned media disrupted network function in a concentration-dependent manner, with unstimulated BV2-conditioned media causing significant disruption (p<0.0193, two-way mixed model, Tukey’s post-hoc test), followed by 10 ng/mL LPS stimulation (p=0.0003) and 100 ng/mL LPS stimulation (p<0.0001), providing further support for NAS as a viable method to quantify complex treatment effects and evaluate disruption of electrophysiological function.

### NAS summarizes neural activity for neurotoxicology screening

Advances in MEA technology [8,19] have led to adoption of MEAs for neurotoxicological screening [20] Given the potential of NAS to consolidate many functional MEA parameters, we sought to determine its applicability to neurotoxicity screening.

For this analysis, NAS values were calculated from MEA toxicity screening of 52 compounds from the NTP or ToxCast libraries [21,22] (Fig 5A-C). In previous studies [21,22], the authors performed a network formation assay (NFA) using primary cortical neurons, measuring 17 parameters of activity in response to compound treatment over 12 days on MEAs to determine compound effects on network formation. Additionally, viability testing was performed to measure cytotoxicity. For each of these assays, EC_50_ values were determined for each compound. Here, we used NAS values to calculate and compare EC_50_ values to individual MEA parameter EC_50_ values and cytotoxicity EC_50_ values (Fig 5D-F, S2 Table).

**Fig 5.**
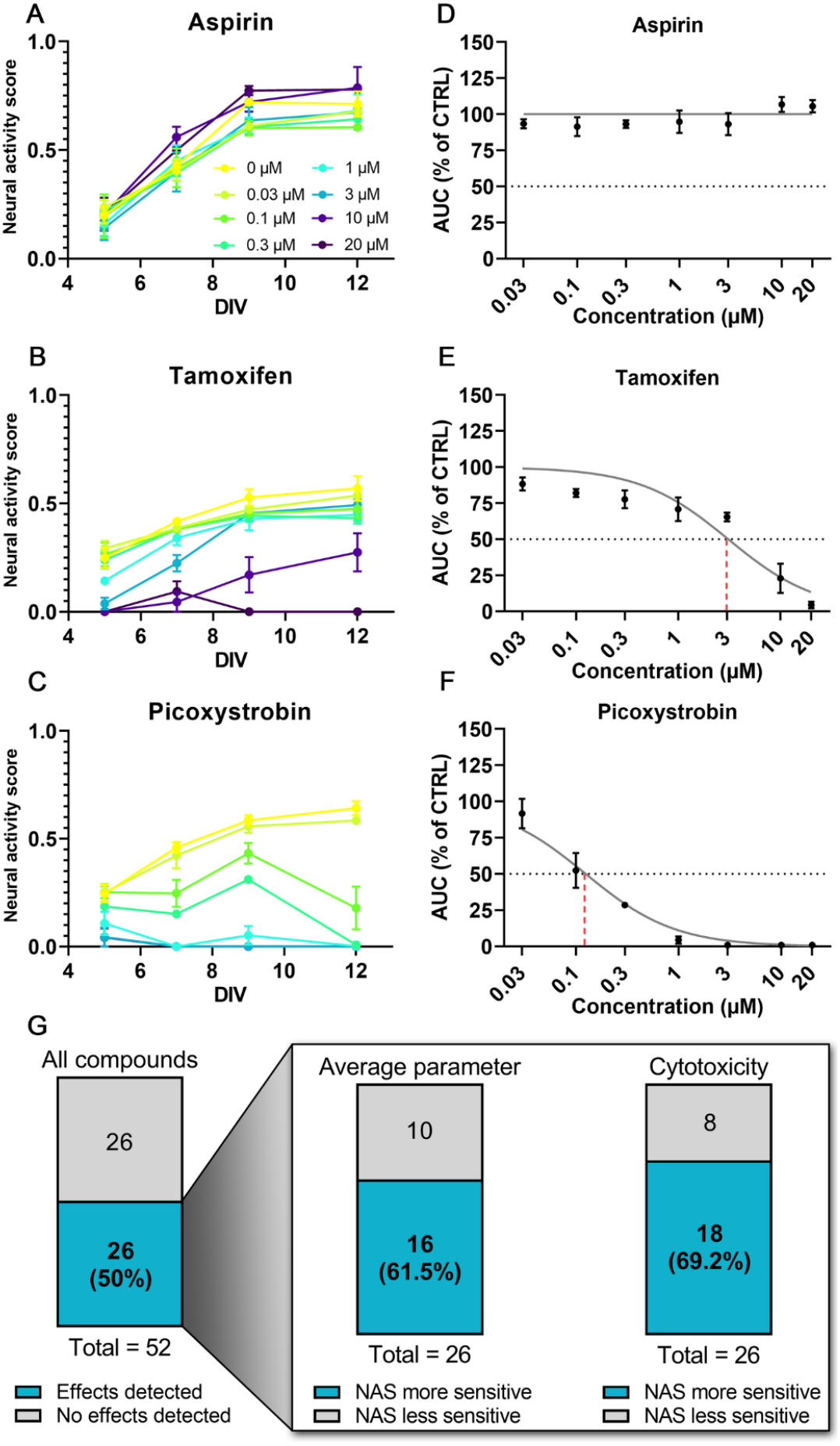
Neural activity score summarizes neural activity for neurotoxicology screening. (A-C) Examples of NAS calculation for all concentrations of three compounds of varying toxicity from EPA compound libraries analyzed. (D-F) Concentration-response curves showing how EC_50_ was determined for the same three compounds. Grey dotted line indicates 50% of control NAS AUC, used as a threshold for EC_50_ extrapolation (indicated via red dashed line). Note the lack of extrapolation for aspirin since sufficient effect was not detected. (G) Summary of NAS EC_50_ values from Frank et al. 2017 [22] and Shafer et al. 2019 [21]. (Left) Total compounds with detected effects (EC_50_ within tested range). (Inset) Sensitivity comparisons for NAS vs. average individual MEA parameter and cytotoxicity assays for all compounds with detected effects. Higher sensitivity is defined as lower EC_50_ value.

Of the 52 compounds we analyzed, 33 were found to have measurable effects in the network formation assay (defined as a decrease in activity by 3X median absolute deviation from control) for at least one activity parameter (though the specific parameter(s) differed among compounds), and 26 compounds were found to have measurable cytotoxicity in the viability assay [21] (Fig 5G). Similarly, using NAS EC_50_ values, we found 26/52 compounds (50%) affected neural activity (Fig 5G). For these compounds, we compared the EC_50_ calculated from NAS to determine sensitivity compared to the average individual MEA parameter EC_50_ values and cytotoxicity EC_50_ values. We found NAS to be more sensitive (lower EC_50_) than the average of all parameters for 16/26 compounds (61.5%) and more sensitive than the average cytotoxicity EC_50_ for 18/26 compounds (69.2%) (Fig 5G, S2 Table).

## Discussion

Advances in MEA technology, including multi-well MEA plates, incubated recording setups, and constantly improving software, allow for higher throughput than previously possible [18,19], though analysis has traditionally been limited to simple parameters, primarily mean firing rate. Only recently have researchers begun incorporating advanced metrics of network activity in these screening approaches [20–22]. These advanced metrics have provided researchers with tools to record from entire neuronal populations and analyze complex neuronal network dynamics. Multi-well MEAs enable high-throughput neuronal recordings, facilitating their adoption for drug/toxicity screening applications and evaluation of complex culture conditions. However, the information that can be gleaned from MEAs has been hampered by limited analytical methods and tools, as well as high variation. Here, we present a novel method to overcome these limitations – development of an index, neural activity score, that incorporates and consolidates traditional MEA measurements into a single quantitative value that can be used to objectively evaluate neuronal network development and function across various culture conditions, treatments, and neural cell sources. This is valuable not only for basic neuroscience research on neuronal networks, but also translational research and preclinical studies.

The results presented here demonstrate the value of NAS to assess potential developmental neurotoxicity (DNT) hazards, a field with a widely recognized need for more sensitive, less variable, and higher throughput functional assays [21–24]. The mixed neural cultures used for NAS derivation and the primary cultures analyzed in the network formation assay are both maturing networks, derived from embryonic stem cells or isolated from neonatal rodents, respectively. As a result, NAS is well-suited for analysis of maturing neural networks, as is necessary in DNT studies, covering a range from non-active to full maturity, with synchronized network bursting. The application to developing networks from multiple cell sources suggest NAS has substantial value for improving the use of MEAs for toxicity screening and drug development.

As the concern over drug development costs continues to rise, scientists are noticing several recurring problems, including the reproducibility crisis and inadequacies of current screening assays, *in vivo* models, and other preclinical studies [25–27]. For neural assays, specifically, assays have traditionally used simple endpoints such as viability and morphological analysis *(i.e.* neurite outgrowth) for screening, primarily due to scalability [28]. However, electrophysiological endpoints are often more sensitive and allow for assessment of electrophysiological toxicity, which involves separate – and highly time-sensitive – mechanisms [8,29,30]. By improving result interpretation, NAS will facilitate incorporation of functional measures into screening programs focused on cytotoxicity and morphology.

Index scoring has been used extensively in clinical settings and *in vivo*; for example, neurological deficits in amyotrophic lateral sclerosis (ALS) and Parkinson’s disease (PD) are assessed using the Revised ALS Functional Rating Scale (ALSFRS-R) [31] and United Parkinson’s Disease Rating Scale (UPDRS) [32], respectively. Stroke severity is frequently measured using various scales *(i.e.* modified Rankin scale (mRS) [33,34] and NIH stroke scale (NIHSS)) [35], and these have been shown to correlate strongly with patient outcomes and be useful for therapeutic evaluation [34]. The simplified analytical pipeline provided by these indices is vital to detecting effects (or lack thereof) in clinical and preclinical studies. Due to this, a need has been recognized to develop multivariate approaches and index scores for *in vitro* approaches, as well [36,37]. Similar analysis pipelines provided by index scores could be especially valuable for screening assays, allowing for improved hit detection when screening potential neurotoxicants or therapeutics. Several composite scores have been developed to condense information from multiple toxicity assays for specific compound classes *(e.g.* endocrine disruptors, halogenated aliphatics), previously [38,39]. Here, we developed NAS using a similar approach to condense the highdimensional data from MEA recordings into a single measurement with reduced variation that can be used to easily and consistently evaluate compound effects on neural activity, as opposed to analyzing many different parameters individually. This reduced variation and improved interpretation could help identify and/or narrow down compounds to examine and develop further, saving time and money wasted on poor candidate compounds. Likewise, improved *in vitro* studies could help reduce the necessity of *in vivo* studies, which are expensive, time-consuming, and have ethical and practical concerns due to a myriad of potential endpoint measurements and species differences that can contribute to high variability and difficulty determining true treatment effects [40].

The validation studies presented here indicate that the NAS formula provides an easily interpretable measure of neural network health/functionality and overall effects of perturbation. By compiling all MEA metrics as opposed to individual metrics (*i.e.* mean firing rate), NAS represents all aspects of neural network function, which can provide more consistent analysis and results interpretation/reporting.

Additionally, NAS has the potential to provide increased sensitivity over a collection of individual parameters, as NAS was more sensitive than the individual parameter average for 61.5% of compounds. This result was interesting, demonstrating the utility of implementing relative parameter weights. Since NAS was derived from how all parameters contribute to development/maturation, this result indicates that this approach may describe treatment effects in a more holistic manner than analysis of individual parameters alone, which only describe certain aspects of activity. However, when specific parameters are of interest, we suggest incorporating NAS as an additional metric for screening, not as a complete replacement, as a summary statistic for electrophysiological function and neural network maturation. Additionally, larger training data sets and/or other optimization may allow for improved sensitivity in the future.

Two of the three compounds for which NAS was found to be most sensitive, MPP+ and picoxystrobin, share similar toxic mechanisms, both inhibiting mitochondrial electron transport chain complexes [41,42]. While further research would be needed to determine if this is more than a coincidence, it does suggest mitochondrial function as a sensitive predictor of neurodegeneration. This finding supports a wealth of evidence linking mitochondrial dysfunction to neurodegenerative diseases, in some cases prior to symptom onset and diagnosis [43–45]. Using NAS to analyze and compare various compound classes in more detail may allow for deeper insight into toxicity mechanisms for different compound classes or varying therapeutic potential in drug discovery studies.

Lastly, challenges in analyzing complex and large data sets have been widely acknowledged across multiple assays and techniques, including high-throughput screening, image analysis, and flow cytometry [46–50]. These challenges include high variability, difficulty interpreting results across multiple metrics, and reproducibility – problems that are only exacerbated when examining complex/emergent phenomena that may be difficult to quantify otherwise, such as neuronal network function. While we developed and validated NAS using MEA data, many of the solutions posed for the aforementioned techniques also utilized PCA and other dimensionality reduction methods, suggesting a similar index scoring approach may be useful for these, and other, applications to gain a deeper understanding of important results.

## Methods

### Cell culture

Mouse HBG3 embryonic stem cell-derived mixed neuronal and glial cells (ArunA Bio, Athens, GA) were cultured according to previously published protocols [11]. Briefly, cells were thawed and seeded on polyethyleneimine (Sigma Aldrich, St. Louis, MO) and laminin (Sigma)-coated MEA plates (Axion Biosystems, Atlanta, GA) in 6 μL droplets centered over the electrode grids at 40-80,000 cells/well. Cells were maintained with media changes every 3-4 days with full neural culture media consisting of BrainPhys™ Basal Media (STEMCELL Technologies, Vancouver, BC, Canada) or Advanced DMEM/F12 (ThermoFisher, Waltham, MA) and AB2 Basal Neural Media (ArunA Bio) (1:1) supplemented with 10% (v/v) KnockOut Serum Replacement (ThermoFisher), 2 mM L-glutamine (ThermoFisher), 1% penicillin/streptomycin (ThermoFisher), 0.1 mM β-mercaptoethanol (Sigma), 10 ng/mL glial-derived neurotrophic factor (GDNF) (Peprotech Inc., Rocky Hill, NJ), and 10 ng/mL ciliary neurotrophic factor (CNTF) (Peprotech).

BV2 microglia cells (gift from Dr. Jae-Kyung Lee, University of Georgia, Athens, GA) were cultured according to previously published protocols [51]. Briefly, cells were thawed and seeded on tissue culture-treated plates at approximately 5-10,000 cells/cm^2^ and passaged at 60-80% confluency. Cells were maintained with media changes every other day with neural medium consisting of DMEM/F12 (ThermoFisher) supplemented with 5% fetal bovine serum (FBS) (GE Healthcare, Chicago, IL), 2 mM L-glutamine (ThermoFisher), and 1% penicillin/streptomycin (ThermoFisher). For lipopolysaccharide (LPS) treatment, cells were treated with 10 ng/mL or 100 ng/mL LPS in neural medium for 24 hours before conditioned media was collected and centrifuged to remove any cells or cellular debris.

### MEA preparation, recording, and data processing

48-well MEA plates (Axion Biosystems) were prepared according to manufacturer’s protocol. Briefly, plates were coated with 0.1% polyethyleneimine (PEI) (Sigma) for 1 hour at 37°C, rinsed with sterile water, and allowed to air dry in a biosafety cabinet overnight. The following day, plates were coated with 20 μg/mL laminin (Sigma) for 2 hours at 37°C prior to cell seeding. Mouse neural cultures (see above) were seeded and allowed to adhere for 1 hour, then maintained in full neural culture media (see above) supplemented with GDNF (Peprotech) and CNTF (Peprotech) (10 ng/mL each) with media changes every 3-4 days throughout the 3-week recording period.

Neuronal activity was recorded using the Maestro system (Axion Biosystems) and AxIS software v2.1-2.5 (Axion Biosystems) with the following settings: band-pass filter (Butterworth, 300-5000Hz), spike detector (adaptive threshold crossing, 8xSD of RMS noise), burst detector (100ms maximum inter-spike interval, 5 spikes minimum, 10 spikes minimum for network bursts, 10ms mean firing rate detection window). Recordings were performed daily for 2 minutes at 37°C after allowing plates to acclimate to the Maestro system.

Raw data files were processed offline using the Statistics Compiler function in AxIS. Statistics Compiler output files were processed in Microsoft Excel (Microsoft Corporation, Redmond, WA) and with custom Python scripts to organize and extract individual parameter data for each well of each MEA plate and for data normalization.

### Neural activity score calculation

After initial data processing, normalization (to z-score values), and outlier removal (−3 > z > 3), JMP 14 (SAS Institute, Cary, NC) was used to conduct principal component analysis. All parameters (S1 Table) were used for all wells (replicates) at 5-19 days *in vitro* (DIV). The first two principal components were used to visually analyze the temporal separation of the data (Fig 2A), then the first principal component was used for linear regression analysis to determine the extent of correlation to time. Finally, the factor loadings for the first principal component were calculated to show the extent of contribution for each individual MEA activity parameter (Table 1).

Factor loadings for principal component 1 were then implemented as coefficients in a formula incorporating, and ultimately consolidating, all of the measured individual MEA parameters into an individual index score – NAS – defined as the sum of each measured parameter value multiplied by its factor loading value for each well (replicate) at each time point (Equation 1).

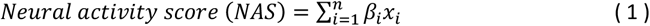

*Equation 1 – where βi are the factor loading values and Xi are the z-normalized measured values for each parameter*

### Analysis of DNT hazard screening data

Raw MEA data (*.raw files generated via the Maestro system and AxIS software (Axion Biosystems), see above) from previous studies [21,22] was processed through the same analysis pipeline described above. Additional processing for neurotoxicity data was based on methods described by Shafer et al. [21], including area under curve (AUC) calculation, Hill function fitting, and EC_50_ extrapolation. Specifically, AUC values for each compound and concentration were calculated in Python 3 using the trapezoidal rule (numpy.trapz() function) to integrate normalized NAS values over time (see Data Availability below for more information about custom Python codes). Concentration-response curves (NAS AUC vs. concentration) were generated via nonlinear least squares regression ([Inhibitor] vs. normalized response model) in GraphPad Prism 8.2.0 (GraphPad Software Inc., San Diego, CA) for each compound with Hill slope = −1.0, and EC_50_ values were extrapolated from the resulting curves.

EC_50_ values corresponding to cytotoxicity (Table S2) that were used for sensitivity analysis were reported from previous studies [21,22]. EC_50_ values for NAS and average MEA parameter were calculated as described above.

### Statistical analysis

Statistical analysis was performed in GraphPad Prism 8.2.0 (GraphPad Software Inc). Two-way repeated measures analysis of variance (ANOVA) was used to assess differences between treatment groups over time for validation studies unless otherwise noted, and post-hoc tests are stated for individual experiments.

## Data availability

Custom Python codes and MEA data (.csv files from AxIS Statistics Compiler, compiled into.xlsx file, and analyzed data at several steps) are provided at the following repository: https://doi.org/10.5281/zenodo.3939310

## Acknowledgments

The authors would like to thank Dr. Tim Shafer and Dr. Katie Paul-Friedman from the Environmental Protection Agency for sharing data and providing advice on toxicological analysis. The authors would also like to thank Dr. Jae-Kyung Lee at the University of Georgia for providing BV2 cells.

## Supporting information

**S1 Fig.**
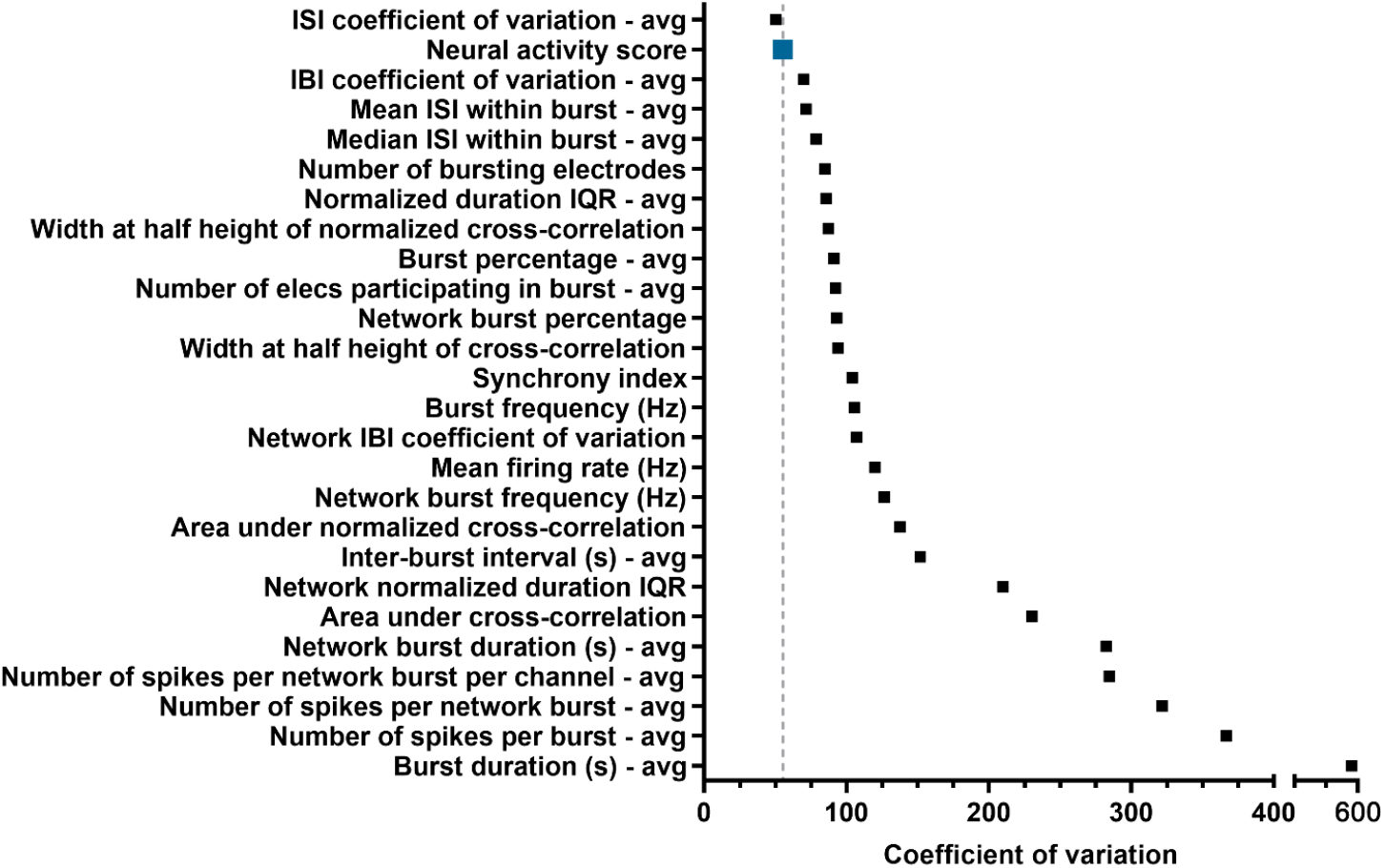
Coefficient of variation values for all individual parameters and neural activity score. Values were calculated from network ontogeny data (Fig 1).

**S1 Table.**
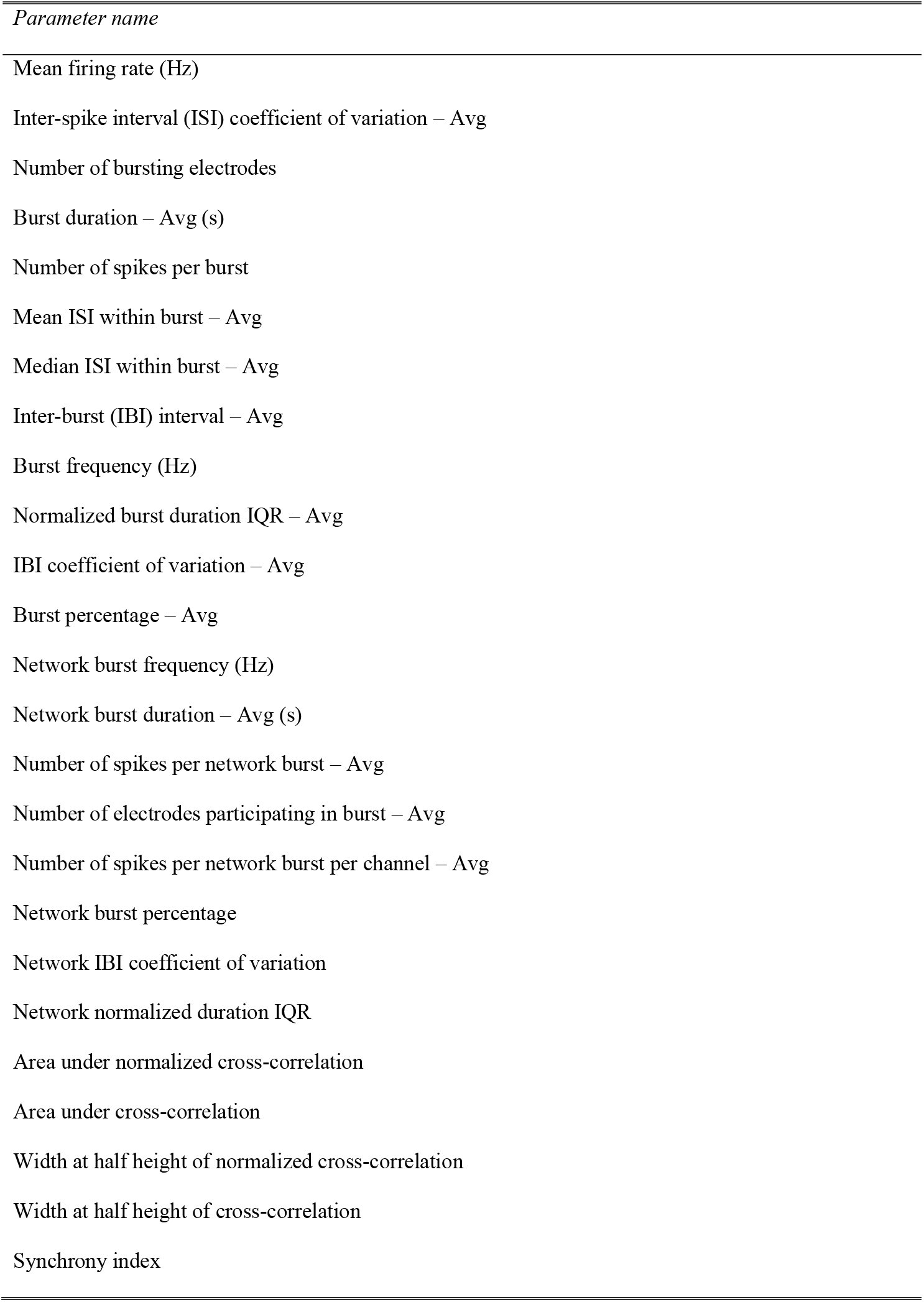
List of all MEA parameters analyzed.

**S2 Table.** EC_50_ values for all compounds analyzed in EPA network formation and toxicity assays. Values were calculated from neural activity score, minimum individual parameter, average of all parameters, and cytotoxicity.

| *Compound* | *CASRN* | *EC<sub>50</sub> (μM)* |  |  |
| --- | --- | --- | --- | --- |
|  |  | *Neural activity score* | *Avg. individual MEA parameter* | *Avg. cytotoxicity* |
| 1-Methyl-4-phenylpyridinium iodide | 36913-39-0 | 4.93 | 4.88 | 6.5483 |
| 1-Ethyl-3-methylimidazolium<br>diethylphosphate | 848641-69-0 | N/A | N/A | N/A |
| 3-Iodo-2-propynyl-N-butylcarbamate | 55406-53-6 | **2.42** | 3.13 | 3.03535 |
| 6 Propyl 2 thiouracil | 51-52-5 | N/A | N/A | N/A |
| Abamectin | 71751-41-2 | **0.16** | 0.37 | 3.9339 |
| Acenaphthene | 83-32-9 | N/A | N/A | N/A |
| Aldrin | 309-00-2 | **4.14** | 4.26 | 7.30305 |
| Aspirin | 50-78-2 | N/A | N/A | N/A |
| Atrazine | 1912-24-9 | N/A | N/A | N/A |
| Auramine O | 2465-27-2 | **2.49** | 3.00 | 3.50965 |
| Benz(a)anthracene | 56-55-3 | N/A | N/A | N/A |
| Berberine chloride | 633-65-8 | **1.83** | 2.03 | 0.2499 |
| Bisphenol AF | 1478-61-1 | N/A | N/A | 16.46775 |
| Bisphenol B | 77-40-7 | N/A | N/A | N/A |
| Boric acid | 10043-35-3 | N/A | N/A | N/A |
| Boscalid | 188425-85-6 | N/A | N/A | N/A |
| Carbamic acid, butyl-, 3-iodo-2-propynyl ester | 55406-53-6 | **1.42** | 1.43 | 1.22425 |
| Chlordane | 57-74-9 | 5.86 | 5.63 | 5.7725 |
| Cloprop | 101-10-0 | N/A | N/A | N/A |
| Clove leaf oil | 8000-34-8 | N/A | N/A | N/A |
| D-Glucitol | 50-70-4 | N/A | N/A | N/A |
| Diphenhydramine hydrochloride | 147-24-0 | 18.12 | 11.56 | N/A |
| Disulfiram | 97-77-8 | 0.85 | 0.64 | 0.0954 |
| Endosulfan | 115-29-7 | **8.71** | 8.84 | 8.90985 |
| Endrin | 72-20-8 | N/A | N/A | N/A |
| Erythromycin | 114-07-8 | N/A | N/A | N/A |
| Estradiol | 50-28-2 | N/A | N/A | N/A |
| Eugenol | 97-53-0 | N/A | N/A | N/A |
| Fenamiphos | 22224-92-6 | N/A | N/A | N/A |
| Fluoxastrobin | 361377-29-9 | **0.69** | 0.89 | 1.1693 |
| Glycerol | 56-81-5 | N/A | N/A | N/A |
| Hexachlorophene | 70-30-4 | 1.92 | 1.83 | 1.8547 |
| Kepone | 143-50-0 | 5.91 | 5.39 | 8.1115 |
| L-Ascorbic acid | 50-81-7 | N/A | N/A | N/A |
| Mancozeb | 8018-01-7 | N/A | N/A | N/A |
| Manganese, tricarbonyl[(1,2,3,4,5-.eta.)-1-methyl-2,4-cyclopentadien-1-yl] | 12108-13-3 | N/A | N/A | N/A |
| Methoxychlor | 72-43-5 | **7.45** | 8.51 | 8.26275 |
| MGK 264 | 113-48-4 | N/A | N/A | N/A |
| Mirex | 2385-85-5 | **3.81** | 4.36 | 2.8233 |
| o,p'-DDT | 789-02-6 | 4.14 | 4.09 | 5.10525 |
| Parathion | 56-38-2 | N/A | N/A | N/A |
| Permethrin | 52645-53-1 | **6.16** | 7.83 | 10.45295 |
| Picoxystrobin | 117428-22-5 | **0.13** | 0.32 | 0.5449 |
| Piperonyl butoxide | 51-03-6 | N/A | N/A | 19.1251 |
| pp-DDD | 72-54-8 | 4.83 | 4.60 | 6.0773 |
| pp-DDE | 72-55-9 | 4.50 | 4.33 | 4.606 |
| pp-DDT | 50-29-3 | 4.36 | 4.33 | 4.0716 |
| Reserpine | 50-55-5 | **1.31** | 1.68 | 9.08055 |
| Rotenone | 83-79-4 | **0.41** | 0.64 | 0.03005 |
| Tamoxifen | 10540-29-1 | **3.09** | 4.35 | 6.4836 |
| Tetracycline | 60-54-8 | N/A | N/A | N/A |
| Triclosan | 3380-34-5 | **9.08** | 9.91 | 9.21815 |

N/A indicates EC<sub>50</sub> not determined within range (0-20 $\mu$ M). Individual parameter averages were only calculated if 13+ (>50%) parameters had determinable ( $0 < EC_{50} < 20$ ) values. Bolded NAS values indicate compounds where NAS was more sensitive (lower EC<sub>50</sub>) than the avg. individual parameter value.

